# *Leishmania amazonensis* infection induces PD-L1 expression on dendritic cells in an mTor-dependent manner and impairs Th1 responses *in vitro* and *in vivo*

**DOI:** 10.1101/2022.02.15.480617

**Authors:** Herbert L. de Matos Guedes, Alessandra M. da Fonseca-Martins, Yuejin Liang, Eric D. Carlsen, Calvin A. Henard, Irina V. Pinchuk, Lynn Soong

## Abstract

*Leishmania amazonensis* is one of the etiological agents of diffuse cutaneous leishmaniasis in South America. In murine models of this infection, dysregulated expansion of effector T cells or exhaustion of Th1 responses are known to be related to pathogenesis, while the induction of regulatory T cells (Tregs) promotes lesion resolution. Recent research has identified several important co-stimulator/receptor pairs, including PD-1/PD-L1, for modulating Th1 responses and Treg induction. In this study, we examined the roles of these molecules in *L. amazonensis*-infected C57BL/6 mice. We found a significant and selective increase in the expression of PD-1 and PD-L1 (20-fold and 5-fold, respectively), in infected footpad tissues, which correlated with an increased percentage of PD-L1^+^CD11c^+^ dendritic cells (DCs) and PD-1^+^CD4^+^ T cells in the draining lymph nodes. To evaluate the mechanism of parasite-induced PD-L1 expression, we infected bone marrow-derived DCs (BMDCs) with promastigotes and amastigotes *in vitro* in the presence of small molecular inhibitors of critical signaling pathways. While *L. amazonensis* infection decreased PD-L2 expression on the surface of BMDCs, infection-mediated PD-L1 up-regulation was consistently detected, which was dependent on mTOR and partially dependent on STAT3, PI3K, and MAPK. Infected BMDCs also significantly inhibited the expansion of Th1 cells, but were more competent in inducing CD25^+^FoxP3^+^ Tregs *in vitro* than the non-infected BMDCs; such effects were PD-L1-dependent, as BMDCs from PD-L1^−/−^ mice failed to do the same. *In vivo* experiments revealed that infected PD-L1^−/−^ mouse tissues showed increased Th1 responses and IFN-γ production, as well as reduced lesion sizes and parasite loads, without affecting Tregs cell expansion. Together, these results support a protective role for PD-1/PD-L1 signaling in regulating local immune responses during *L. amazonensis* infection. This study provides new insights on immune regulation in New World cutaneous leishmaniasis.

## Introduction

Leishmaniasis is a parasitic neglected tropical disease caused by multiple *Leishmania* spp. Clinical manifestations of the disease range from localized cutaneous lesions to lethal visceral infection, and disease severity and responsiveness to treatment vary widely depending on the infecting parasite species and the host immune status (1). Diffuse cutaneous leishmaniasis is a rare form of the disease characterized by multiple, chronic, heavily-parasitized cutaneous lesions, T cell anergy to parasite antigens, and refractoriness to therapy (2).

In mice, *L. amazonensis* infection does not obey the strict patterns of Th1/Th2 polarization that are classically observed in murine *L. major* infection (3). BALB/c and C57BL/6 mouse strains each generate a mixed Th1/Th2 cell and cytokine pattern in response to *L. amazonensis* infection, reminiscent of leishmaniasis in humans (3-5). Of interest, transgenic mice lacking RAG2 (Rag2^−/−^) and major histocompatibility complex class II (MHC II^−/−^) are resistant to *L. amazonensis* infection, suggesting a pathological role for effector CD4^+^ T cells (3).

Both *L. major* and *L. amazonensis* induce expansion of regulatory T cells (Tregs), but these cells appear to play opposing roles in promoting parasite clearance. In *L. major* infection, Tregs are associated with parasite persistence (6), while Tregs during *L. amazonensis* infection temper inflammation and facilitate disease resolution (7). The mechanism through which Tregs are induced in *L. amazonensis* infection, and how this process could potentially be amplified, remains poorly characterized at this time.

The PD-1/PD-L1 signaling pathway is critically important in crosstalk between T cells and antigen presenting cells (APC) and in the induction of T cell anergy, a phenomenon that has been documented in both the organ transplant setting (8) and in different types of tumors (9). In murine infection by *L. mexicana*, it was shown that PD-L1^−/−^ mice had smaller lesions and lower parasite loads when compared to the background C57BL/6 mice (10). We have previously shown that dendritic cells (DCs) in the draining lymph nodes of *L. amazonensis*-infected BALB/c mice express more PD-L1 (11), and that treatment of infected mice *in vivo* with anti-PD-1 and anti-PD-L1 monoclonal antibodies promotes T cell reinvigoration and control of parasite replication. However, these data were generated using a susceptible mouse strain, and the role of PD-1/PD-L1 in a more resistant genetic background (e.g. C57BL/6 mice), as well as the specific cell interactions and downstream effects that contribute to this phenomenon, remain to be elucidated.

DCs, natural killer (NK) cells, and macrophages are suppressed by *L. amazonensis* amastigotes infection. Pathways involving mitogen-activated protein kinases (MAPK) and extracellular signal-regulated protein kinases (ERK) have been implicated in parasite-driven impairment of CD40 expression in DCs (12, 13), while in infected macrophages, phosphoinositide 3-kinase (PI3K) and protein kinase B (Akt) have been linked to impaired IL-12 production (14). Furthermore, the production of TGF-β has been associated with the T cell anergy observed during *L. amazonensis* infection (15). There are molecules involved in DC suppression and the capacity to induce T cell anergy and Tregs, as the activation of STAT3 that promotes PD-L1 expression.

STAT3 transcription factor signaling promotes suppression of DCs, inhibiting production of inflammatory cytokines/activator markers and promoting tolerance (16). Both tumor cells and the bacterium, *Helicobacter pylori*, have been reported to activate STAT3, impairing DC maturation (17). The process of STAT3 activation involves different pathways: Janus kinases, the MAPK pathway, and the PI3K pathway. The major mechanism induced by STAT3 signaling to promote tolerance and Tregs is the induction of PD-L1 on DCs, in this case denominated as tolerogenic APCs (18). PD-L1 is one of the most important co-stimulatory molecules involved in the induction of T cell anergy (19) and the induction, function, and maintenance of Tregs (20). *Mycobacterium tuberculosis* (21), *Staphylococcus aureus* (22), and lung microbiota (23) induce Tregs via PD-L1/PD-1, indicating that this is a conserved strategy among microorganisms to modulate the host immune response. Based on the capacity of *L. amazonensis* to suppress DCs and induce MAPK-ERK (13) and PI3K (24), here we investigate the hypothesis that *L. amazonensis* infection induces PD-L1 expression on DCs in order to induce Tregs and inhibit T-bet via the PD-1/PD-L1 interaction.

In this work, we have used both bone marrow-derived DCs (BMDCs) *in vitro* and murine models to confirm the role of PD-L1/PD-1 in inhibiting Th1 responses during *L. amazonensis* infection and the dependence of this phenomenon on mTOR in DCs.

## Materials and Methods

### Mice

Female C57BL/6, BALB/c (Taconic), *Myd88*^−/−^, PD-L1^−/−^ and MHCII^−/−^ mice (both on the C57BL/6 background, obtained from the University of Texas Medical Branch) were used in this study. Mice were maintained under specific pathogen-free conditions and used for infection experimentation at 6-8 weeks old, according to protocols approved by the Institutional Animal Care and Use Committees.

### Parasite culture

Infectivity of *L. amazonensis* (strain RAT/BA/74/LV78) was maintained by regular passage through BALB/c mice. Promastigotes were cultured at 26°C in Schneider’s *Drosophila* medium (Invitrogen, Carlsbad, CA, USA), pH 7.0, supplemented with 20% fetal bovine serum (FBS; Sigma, St. Louis, MO, USA), 2 mM L-glutamine, and 50 μg/ml gentamicin. Cultures of stationary-phase promastigotes of less than five passages were used for DC or animal infection and for the generation of axenic amastigotes. Axenic amastigotes were generated by culturing stationary-phase promastigotes at 32°C in Grace’s insect cell culture medium (Invitrogen), pH 5.2, supplemented with 20% heat-inactivated FBS and 25 μg/ml gentamicin. The amastigotes were used for infections after the third passage.

### *In vivo* infection

C57BL/6 mice (5/group) were infected subcutaneously in the right hind footpad with 5×10^6^ stationary-phase promastigotes of *L. amazonensis*. Lesion size was monitored with digital calipers (Control Company, Friendswood, TX, USA), and parasite burdens were measured via limiting dilution assay as previously described (25). After 9-12 weeks, popliteal draining lymph nodes (LNs) were collected from individual mice, macerated with a tissue mixer and the cells were stained immediately for the expression of markers by flow cytometry. The footpad tissue was collected and macerated for real-time PCR assay and the supernatants collected and cytokines quantified by individually assaying for the presence of IL-4, IFN-γ and IL-17 by specific ELISAs using a standard protocol, detection limits for IL-4 is 7.8–500 pg/ml (BD OptEIA, San Diego, USA), IFN-γ is 9.4 – 600 pg/mL (R&D Systems, Minneapolis, USA) and IL-17 is 10.9 – 700 pg/mL (R&D Systems, Minneapolis, USA).

### Quantitative reverse transcriptase PCR (qRT-PCR)

Total RNA was extracted from footpad tissues by using the RNeasy Mini kit (Qiagen, Valencia, CA, USA) and digested with RNase-free DNase (Qiagen). cDNA was synthesized with the iScript cDNA Synthesis kit (Bio-Rad Laboratories, Hercules, CA, USA). The abundance of target gene expression was measured by qRT-PCR using a Bio-Rad CFX96 real-time PCR apparatus and a SYBR Green Master Mix (Bio-Rad) for all PCR reactions. PCR reactions were started at 95°C for 3 min, followed by 39 cycles of 95°C for 10 sec and 60°C for 10 sec, and ended with an elongation step at 72°C for 10 sec. Dissociation melting curves were obtained after each reaction to confirm the purity of PCR products. Relative abundance of mRNA expression was calculated by using the 2^−ΔΔCT^ method. Actin was used as the housekeeping gene for all analyses of the footpad tissue. Primer sequences are listed in Table S1.

### DC generation and *in vitro* infection

Bone marrow cells were flushed from the femur and tibia of BALB/c or C57BL/6, *Myd88*^−/−^ or PD-L1^−/−^ mice with RPMI/10% FBS and then were centrifuged at 400 g for 7 min at 24°C. BMDCs were grown in complete RPMI 1640 (Sigma-Aldrich) containing 10% FBS, supplemented with 20 ng/ml rGM-CSF (eBioscience, San Diego, CA, USA). On day 8, DCs were harvested and adjusted to 5×10^5^/well in 48-well plates. After 8-9 days, cells were incubated with carboxyfluorescein succinimidyl ester (CFSE)-labeled parasites (5:1 parasite-to-cell ratio for amastigotes, or 10:1 for promastigotes; staining with CFSE (0.5 μM,- Invitrogen) at 37°C. At 24 h post-infection, cells were collected for flow cytometry analysis (CD11c^+^, PD-L1^+^, PD-L2^+^), and the supernatants were harvested for cytokine detection (IL-10 and TGF-β) by specific ELISAs using a standard protocol (BD OptEIA). The detection limits for IL-10 is 31.3–2000 pg/ml e TGF- β is 62.5–4,000 pg/mL). For the study of the intracellular mechanisms involved in PD-L1 induction, cells were treated with a STAT3 inhibitor (150 nM), STAT5 inhibitor (10 uM), rapamycin (100 ng/ml), wortmannin (50 nM), U0126, MEK inhibitor (100 nM), SB 203580, MAPK inhibitor (100 nM), Janus kinase I inhibitor (10 nM), Janus kinase II inhibitor (10 nM), or Janus kinase III inhibitor (10 nM), all from Sigma-Aldrich, 1 h prior to the infection. Controls were performed with medium or DMSO as appropriate. For the determination of percentage of PD-L1 inhibition, we considered the increase of PD-L1 expression on infected BMDCs in comparison to the non-infected (medium) control as 100% of activity. The reduction of PD-L1 expression on treated on BMDCs was then evaluated using the ratio of BMDC-infected to BMDC-medium and the percentage of inhibition was determined by comparing these values in relation to the controls.

### Cell Staining for Flow Cytometry

Cells from lymph nodes (1×10^6^) or DC *in vitro* culture (5×10^5^) were washed with PBS at 400 g for 5 min at 4°C and blocked with FcX (BioLegend) for 15 min, followed by staining with the antibody cocktail for 30 min at 4°C. Cells were then washed with a cytometry buffer (PBS with 5% FBS) at 400 g for 5 min and 4°C, then fixed with 4% formaldehyde (Sigma) for 15 min at 4°C. Cells were washed and resuspended in the cytometry buffer and stored in the dark at 4°C until acquisition. After that, we used the fixable Live/Dead dye (efluor506) and the following antibodies were used: anti-CD11c (PerCP-Cy5.5), PD-L1 (APC) and PD-L2 (PE), MHC II (FITC), CD4 (PE), CD8 (APC-Cy7), PD-1 (FITC), FoxP3 (FITC), CD25 (PerCP-Cy5.5), T-bet (PE-Cy7), IFN-γ (APC) (eBioscience). Acquisition of events (100,000 events) was performed on a LSR BD Fortessa. The gate strategy was performed based on the selection of cell size (FSC) and composition (SSC). After identifying the main population, a gate of FSC-A (area) and FSC-H (weight) was used, where cellular doublets were excluded. Gates for positive events were established through Fluorescence Minus One (FMO) control. The data analyzes were performed using the FlowJo software (Oregon, USA).

### Western immunoblotting

BMDC from C57BL/6 mice were infected with amastigotes (ratio 5:1) of *L. amazonensis,* overnight. Cells were washed in PBS and cell pellets were lysed using CelLytic M (Sigma) supplemented with complete Mini EDTA-free protease inhibitor cocktail (Roche Applied Science, Indianapolis, IN, USA). Cellular debris was removed by centrifugation at 13,000 × *g*. Protein was quantified, 20 μg was loaded onto 10% Tris-Glycine gels and separated using the XCell SureLock Blot Module (Invitrogen). Gels were transferred to polyvinylidene fluoride (PVDF) membranes. Membranes were blotted using the primary antibodies: Ebi3 (2.5 μg/ml; eBioscience) or IL-12/IL-35 p35 (1 μg/ml; R&D Systems), phospho-mTOR (Ser2448, Invitrogen), mTOR (7C10, Invitrogen), and Glyceraldehyde-3-phosphate dehydrogenase (GAPDH) (Novus Biologicals, Littleton, CO, USA). Primary antibodies were followed by HRP-conjugated secondary antibodies (Jackson ImmunoResearch Laboratories, West Grove, PA, USA). Protein was visualized using the SuperSignal West Pico ECL kit (Pierce – Thermo Fisher Scientific, Rockford, IL, USA).

### DC: T cell co-culture

Anti-CD3 was used to coat the wells of a 48-well plate (1 μg/well) overnight. BMDCs (5 × 10^5^) from C57BL/6 and PD-L1^−/−^ mice, generated as previously mentioned, were plated in the presence or absence of *L amazonensis* amastigotes (ratio 5:1) or not for 18 h. CD4^+^CD25^−^ T cells were purified from spleens of C57BL/6 mice by negative selection using the EasySep™ Mouse Naïve CD4^+^ T cell Isolation Kit following the manufacturer’s instructions. Naive CD4^+^ T cells (5 × 10^5^) were co-cultured with BMDCs in complete RPMI 1640 (Sigma-Aldrich) containing 10% FBS and IL-2 (1 ng/ml). After 5 days, cells were analyzed by flow cytometry (T-bet and FoxP3 on CD4^+^ T cells).

### Data analysis

Results are expressed as mean ± SEM or SD with a confidence level of p ≤ 0.05. For multiple comparisons, a one-way ANOVA followed by Tukey pairing was performed. Cell analyses were performed using the paired Student’s t-test (p values are indicated in the graphs and figure legends). Data analysis was performed using GraphPad Prism^®^ 8.00 software (San Diego, California, USA).

## Results

### *L. amazonensis* infection induces PD-1 and PD-L1 *in vivo*

Four major receptor-ligand pairs have been associated with the co-stimulatory response between APCs and T cells: PD-1/PD-L1, ICos/ICosL, OX-40/OX-40L, and GITR/GITRL. To evaluate which pairs were enhanced during *L. amazonensis* infection, we compared transcript levels of these molecules in naïve and parasite-infected footpad tissue via real-time PCR. We observed 20 fold greater expression of PD-1 (Fig. 1A) and 5 fold greater expression of PD-L1 (Fig. 1E) in the parasite-infected tissues of C57BL/6 mice, while levels of the other molecule pairs were minimally altered or not were altered by infection (Fig. 1 and Suppl. Fig.1).

**Fig 1.**
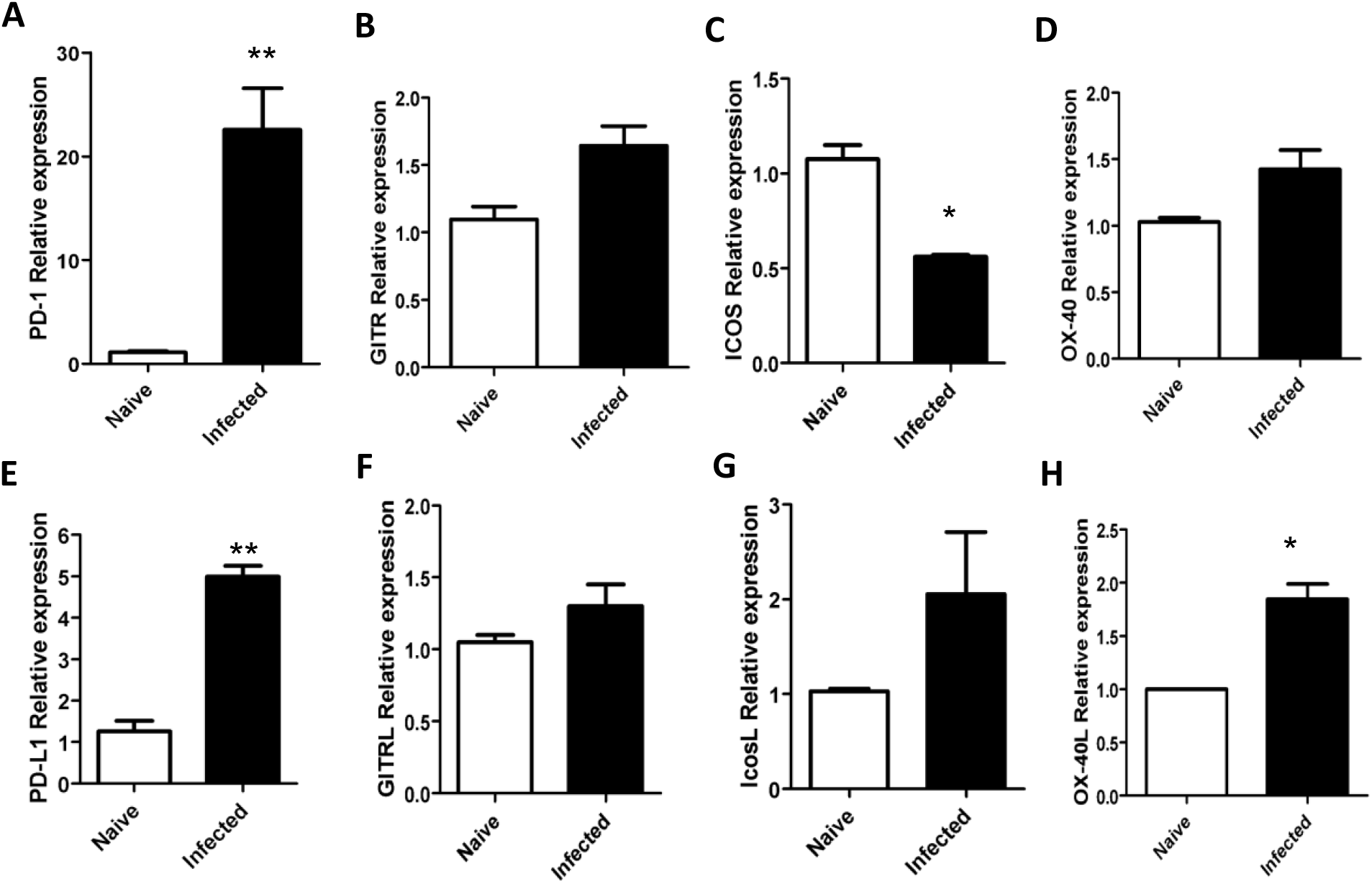
*Leishmania amazonensis* induces PD-1 and PD-L1 expression in the infected footpads of mice. C57BL/6 mice were infected with 5×10^6^ promastigotes in the hind footpad. Naive= not infected. After 3 months, infected footpads were evaluated by real-time PCR for PD-1 (A), PD-L1 (E), GITR (B), GITRL (F), ICos (C), ICosL (G), OX40 (D), and OX40L (H). Actin was used as a housekeeping gene. Data ± SD of individual mice (5 mice/group) are representative of three independent experiments producing the same result profile. *P<0.05, **P<0.01.

We further evaluated PD-1 and PD-L1 expression in *L. amazonensis* infection *in vivo* by quantifying positive cell populations via flow cytometry. We found that the number and percentage of PD-1-expressing CD3^+^CD4^+^ T cells increased roughly 5-fold in response to infection (Fig. 2, and Suppl. Fig.1), and that PD-L1-expressing CD11c^+^ DCs also significantly increased in percentage and number in MHCII^hi^ (Fig. 3, and Suppl. Fig.1). We did not observe a difference in the expression of PD-1 on CD3^+^CD8^+^ T cells between naïve and infected mice (data not shown).

**Fig 2.**
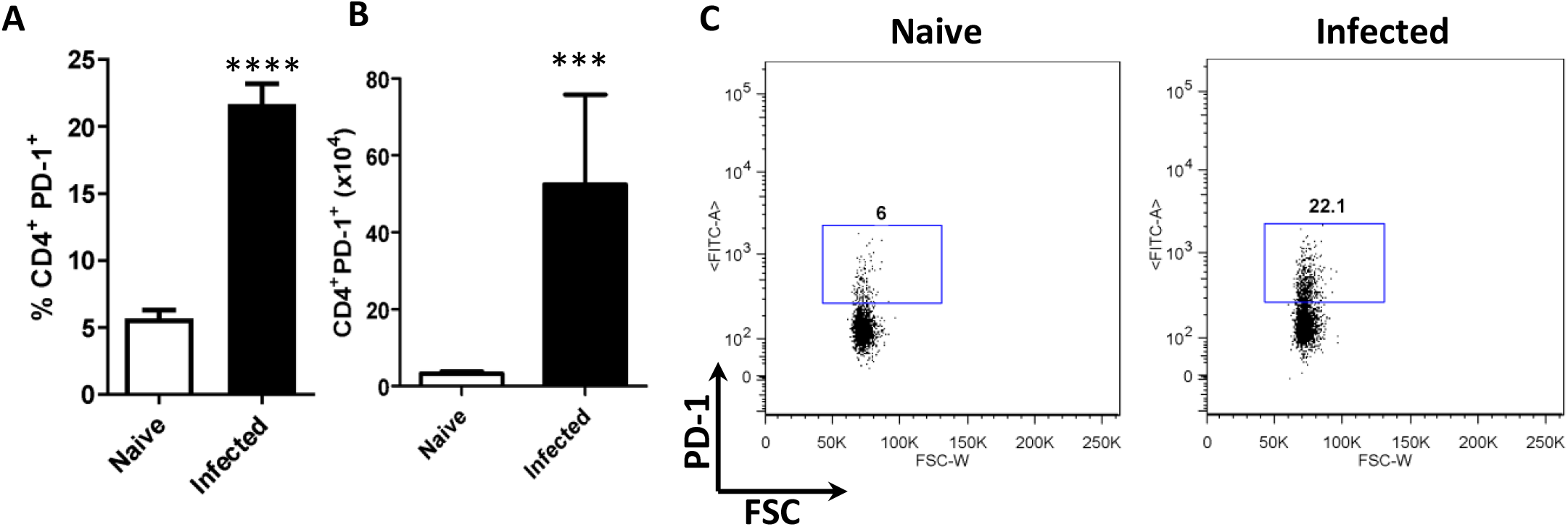
*Leishmania amazonensis* infection induces PD-1 on CD4^+^ T cells *in vivo*. C57BL/6 mice were infected with 5×10^6^ promastigotes in the hind footpad. Naive= not infected. After 3 months, the cell populations of the popliteal draining lymph nodes were evaluated by flow cytometry. A) Percentage of CD4^+^PD-1^+^ T cells. B) Number of CD4^+^PD-1^+^ T cells. C) Dot plot of PD-1 expression on CD4^+^ T cells (PD-1-FITC, FSC-cell volume). Naïve = not infected. Data ± SEM of individual mice (5 mice/group) are representative of three independent experiments producing the same result profile. **P<0.01, ***P<0.001.

**Fig 3.**
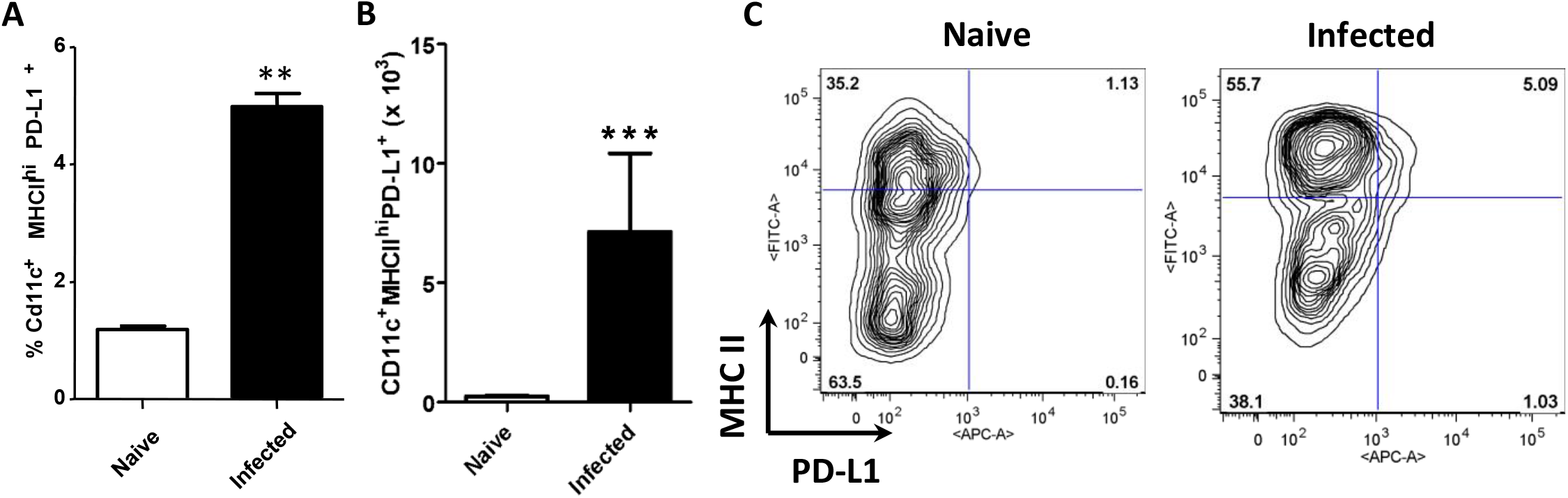
*L. amazonensis* infection *in vivo* induces expression of PD-L1 on DCs. C57BL/6 mice were infected with 5×10^6^ promastigotes in the hind footpad. Naive= not infected. After 3 months, the cell population of the popliteal draining lymph nodes were evaluated by flow cytometry. A) Percentage of CD11c^+^ MHCII^hi^ PD-L1^+^ cells. B) Number of CD11c^+^ MHCII^hi^ PD-L1^+^ cells. C) Dot plot of MHCII and PD-L1 expression (MHCII-FITC, PD-L1-APC) on the CD11c^+^ cells. Naïve = not infected. Data ± SD of individual mice (5 mice/group) are representative of three independent experiments producing the same result profile. * P<0.05, **P<0.01, ***P<0.001.

### *L. amazonensis* induces PD-L1, but inhibits PD-L2, expression in dendritic cells *in vitro*

We evaluated *in vitro* the capacity of *L. amazonensis* to induce PD-L1 without any extra stimulus, which would be indicative of the parasite being able to modulate DCs by an innate mechanism. We evaluated PD-L1 and PD-L2 expression after both promastigote and amastigote infection of BMDCs from C57BL/6 mice (Fig. 4). CFSE-labeled parasites were used to distinguish infected cells from bystander cells. We observed that parasite-carrying BMDCs (infected with either amastigotes or promastigotes) had significantly increased expression of PD-L1 (Fig. 4B and D) and reduced expression of PD-L2 when compared to the uninfected bystander cells (Fig. 4C). Similar results were also observed using BMDCs from BALB/c mice (Suppl. Fig. 2). The ability of the parasite to induce PD-L1 in DCs could be an explanation as to how *L. amazonensis* induces suppression of T cell immune response.

**Fig 4.**
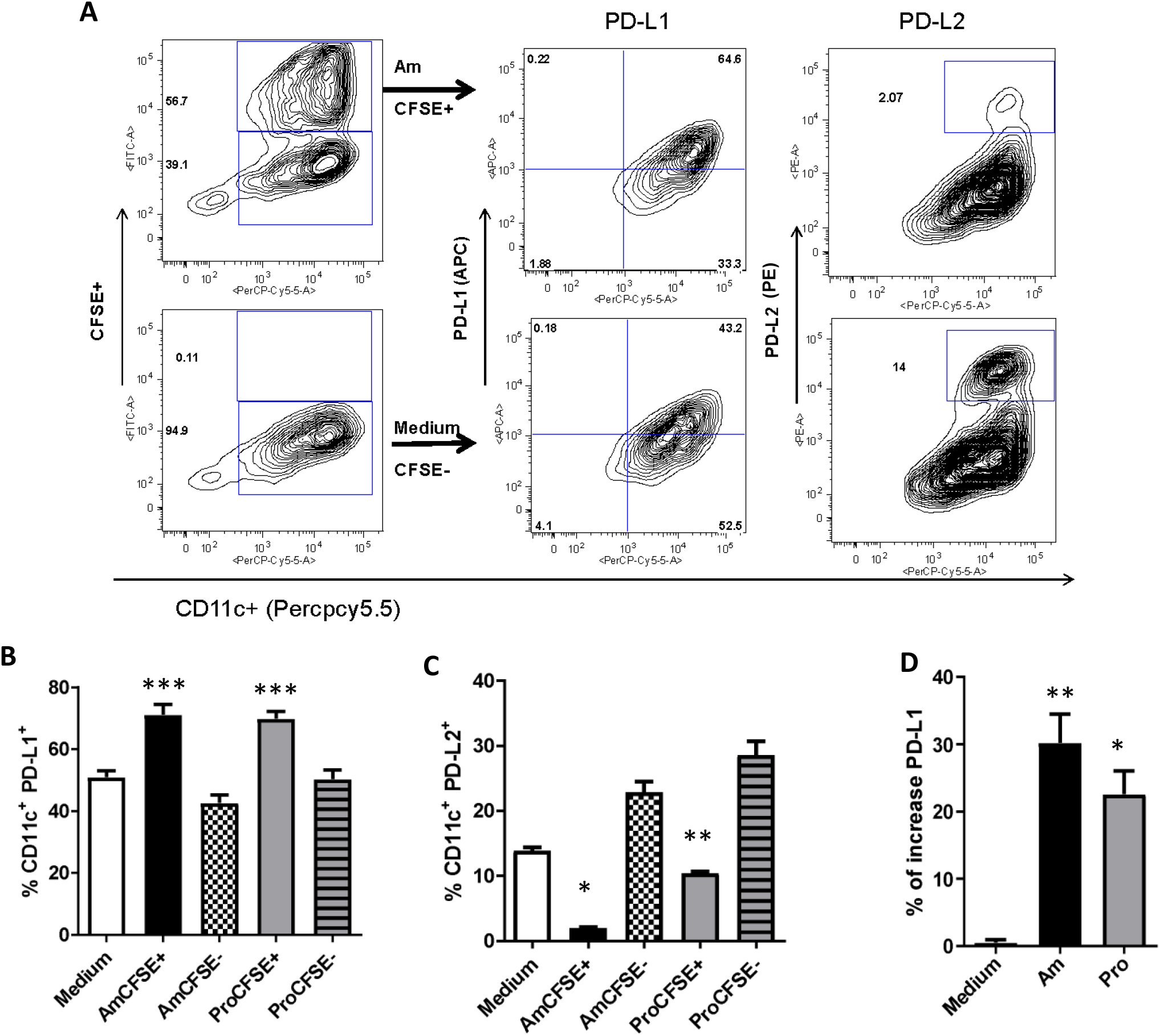
Induction of PD-L1 expression and inhibition of PD-L2 expression on infected dendritic cells. BMDCs (5×10^5^) from C57BL/6 mice were infected for 18 h with *L. amazonensis* amastigotes (Am, ratio 5:1) or promastigotes (Pro, ratio 10:1). Medium= not infected cells. Cells were analyzed by flow cytometry for CFSE^+^ (*L. amazonensis*), CD11c^+^ (PerCP-Cy5.5), PD-L1^+^ (APC), and PD-L2^+^ (PE). A) Dot plot of CFSE^+^CD11c^+^ cells and CFSE^−^CD11c^+^ cells, followed by the expression of PD-L1^+^ and PD-L2^+^ cells on the CD11c^+^ cells. B) Percentage of CD11c^+^PD-L1^+^ cells. C) Percentage of CD11c^+^PD-L2^+^ cells. D) Percentage increase of PD-L1 expression. Data ± SEM are representative of three independent experiments producing the same result profile. * P<0.05, **P<0.01, ***P<0.001.

### *L. amazonensis* infection induces PD-L1 expression by the mTOR pathway

Our next step was to evaluate the PD-L1 induction mechanism in infected DCs. There are two major signaling pathways that can lead to PD-L1 expression: mTOR–PI3K–STAT3 and MAPK-ERK-STAT3 (which may or may not include utilization of Janus kinases). Since *L. amazonensis* has already been shown to induce PI3K and MAPK/ERK, we suspected that these pathways may be involved in PD-L1 induction. Using different inhibitors, we evaluated the pathways involved in PD-L1 induction in DCs (Fig. 5). When infected BMDCs were treated with the PI3K inhibitor, wortmannin (Fig. 5A) and the STAT3 inhibitor (Fig. 5B), approximately 40% of parasite-induced PD-L1 expression was inhibited. However, when the mTOR inhibitor, rapamycin, was used (Fig. 5A), 60% inhibition occurred, demonstrating that the major mechanism of PD-L1 induction in DCs is dependent on mTOR. STAT5 inhibition resulted in an increase in PD-L1 expression on both infected and bystander BMDCs, suggesting that STAT5 signaling acts to dampen PD-L1 induction (Fig. 5B). Using Janus kinase inhibitors (JKI, JKII, and JKIII), we could not detect any modulation in parasite-induced PD-L1 expression, implying no participation of cytokines in PD-L1 induction by *L. amazonensis in vitro* (Fig. 5C). MAPK inhibition (SB 203580) (Fig. 5D), MEK1/2 inhibition (UO126) and the use of *Myd88*^−/−^ BMDCs (Fig. 5E) produced a small inhibition of PD-L1. A comparison of the inhibitor effect of each of these treatments on parasite-induced PD-L1 expression is summarized in Fig. 5F. Since PD-L1 induction appeared to be dependent on mTOR, we next assessed whether parasites could induce p-mTOR. For this, Western blot analysis was performed on *L. amazonensis*-infected DCs, which indicated p-mTOR activation (Suppl. Fig.3). These results suggest a major participation of mTOR, with contribution of PI3K and STAT3, in the induction of PD-L1 expression by *L. amazonensis*.

**Fig 5.**
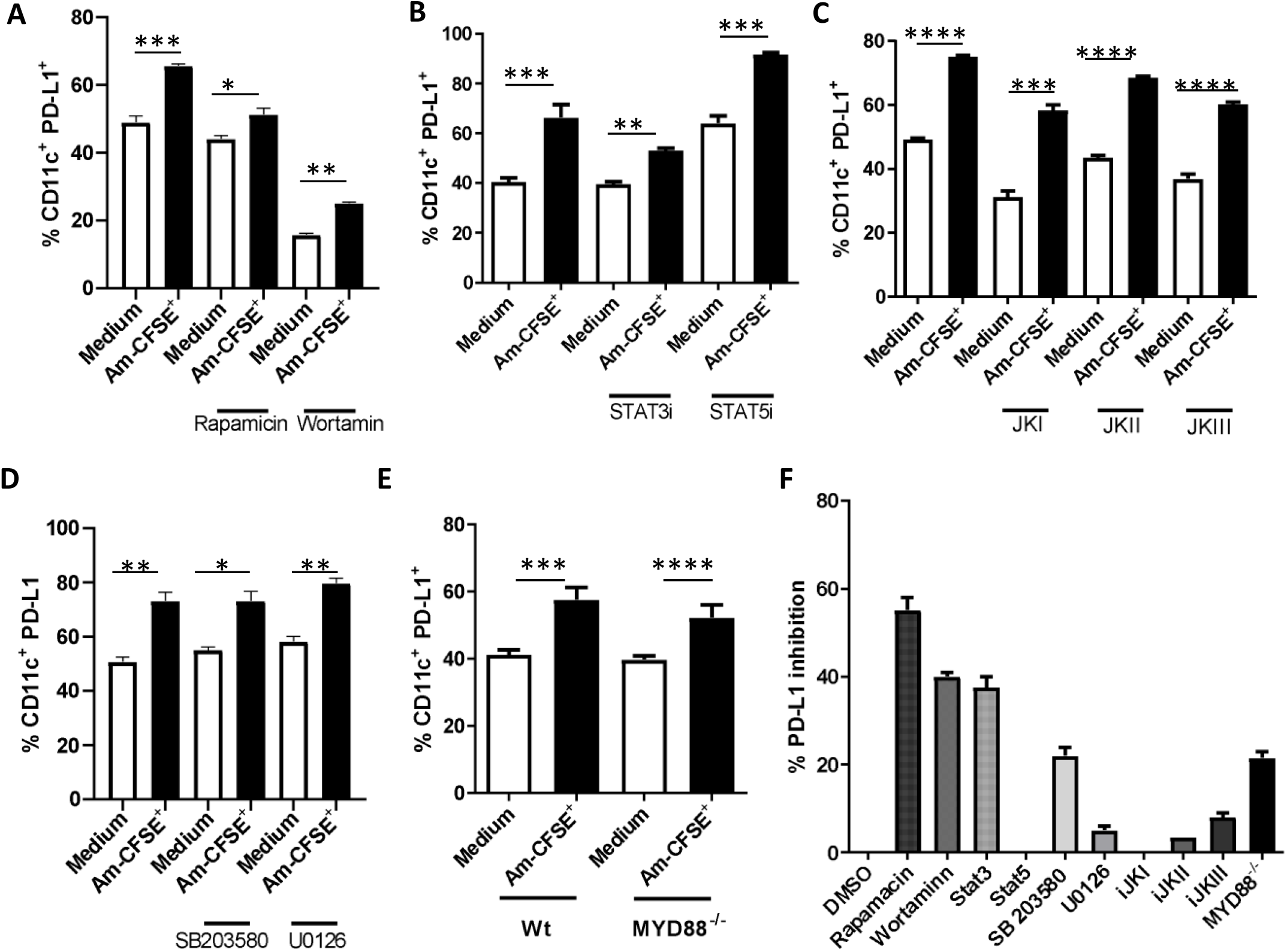
*L. amazonensis* infected DC induces PD-L1 expression. BMDCs (5×10^5^) from C57BL/6 and PD-L1^−/−^ mice were treated with inhibitors (where indicated), or maintained in medium with DMSO as a control, for 1 h, after which, cells were infected with CFSE-labeled *L. amazonensis* amastigotes (ratio 5:1) for 24 h. Medium = non-infected cells. Analysis of the cell markers was performed by flow cytometry. A) Percentage of CD11c^+^PD-L1^+^ cells following incubation with rapamycin (100 ng/ml) and wortmannin (50 nM). B) Percentage of CD11c^+^PD-L1^+^ cells following incubation with a STAT3 inhibitor (150 nM) and a STAT5 inhibitor (10 μM). C) Percentage of CD11c^+^PD-L1^+^cells following incubation with the Janus kinase inhibitors: JKI (10 nM), JKII (10 nM), and JKIII (10 nM). D) Percentage of CD11c^+^PD-L1^+^cells following incubation with U0126, MEK inhibitor (100 nM) and SB 203580, MAPK inhibitor (100nM). E) Percentage of CD11c^+^PD-L1^+^ cells following infection of BMDCs from C57BL/6 and *Myd88*^−/−^ mice. F) Percentage of PD-L1 inhibition on CD11c^+^ cells. Data ± SEM are representative of four independent experiments producing the same result profile. * P<0.05, **P<0.01, ***P<0.001, ****P<0.0001.

### *L. amazonensis* infected DCs inhibit generation of Th1 T cells

To evaluate the role of PD-L1 induction in infected DCs on T cell differentiation *in vitro*, we performed co-cultures using BMDCs from C57BL/6 or PD-L1^−/−^ mice and naïve CD4^+^ T cells from C57BL/6 mice. After 5 days, we observed a small increase in the percentage of CD4^+^CD25^+^FoxP3^+^ cells in the co-culture using infected BMDCs from C57BL/6 mice in comparison to non-infected BMDCs from the same mice (medium), as well as a substantial reduction in the percentage of T-bet^+^ T cells in infected co-cultures. This inhibition of T-bet was not observed when the T cells were co-cultured with BMDCs from PD-L1^−/−^ mice, indicating the role of PD-L1 in the inhibition of effector cell induction (Fig. 6C and D). Furthermore, infected BMDCs from PD-L1^−/−^ mice failed to promote an increase in the Treg proliferation in relation to the non-infected BMDCs from PD-L1^−/−^ mice (Fig. 6A and B). This result indicates a role of PD-L1 in the inhibition of effector T cells and Treg induction by *L. amazonensis* infection. We could not detect any increase in the supernatant levels of TGF-β and IL-10 from BMDC culture following infection indicating no modulation in regulatory/suppressive cytokines (Suppl. Fig. 4).

**Fig 6.**
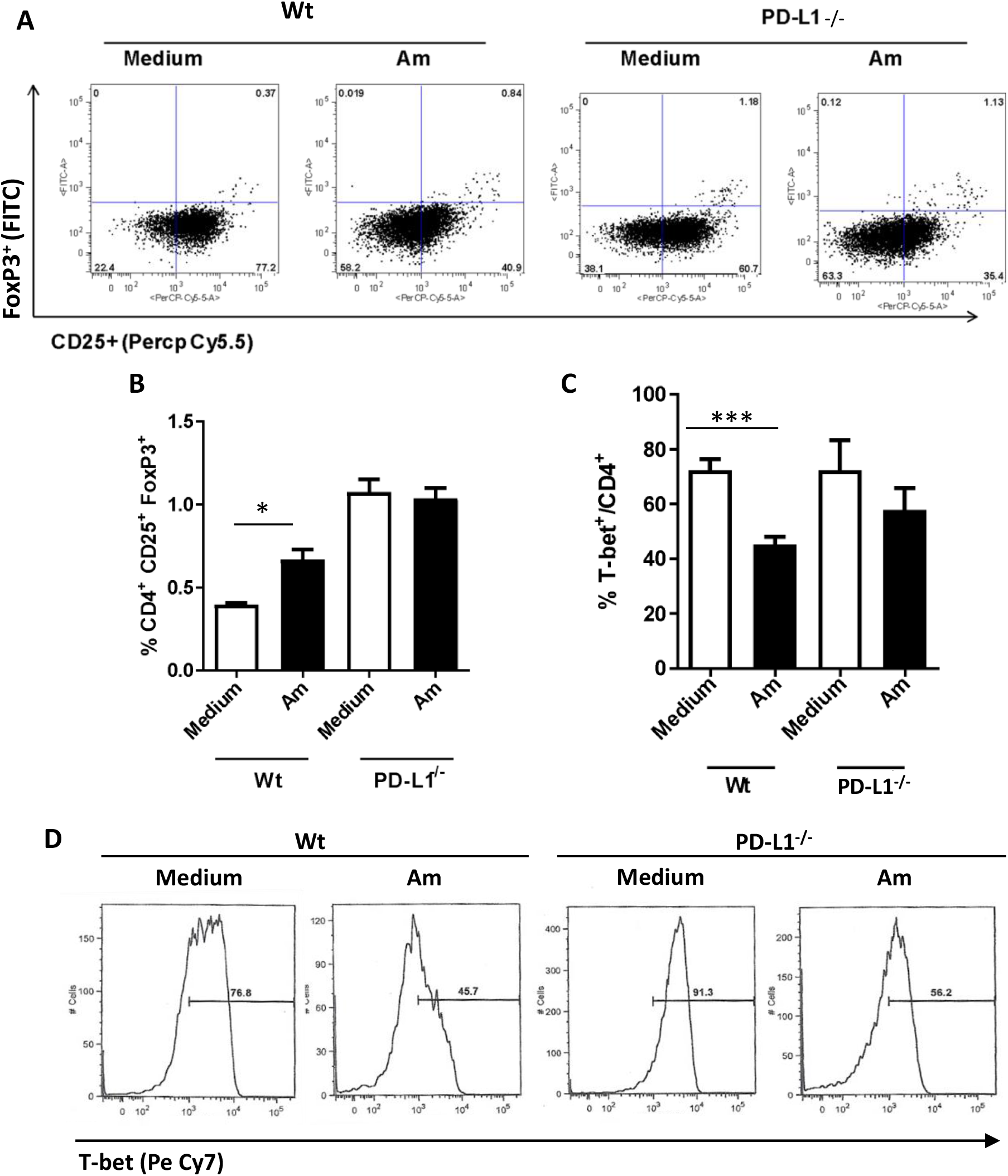
Dendritic cells from PD-L1^−/−^ mice fail to promote Treg proliferation. BMDCs (5×10^5^) from C57BL/6 (Wt) and PD-L1^−/−^ mice were infected with *L. amazonensis* amastigotes (Am; ratio 5:1), and then co-cultured with naïve CD4^+^ T cells from C57BL/6 mice. After 5 days, cells were analyzed by flow cytometry. Medium = non-infected cells. A) Dot plot of CD4^+^CD25^+^FoxP3^+^ cells (FoxP3^+^-FITC and CD25^+^-PerCP-Cy5.5). B) Percentage of CD4^+^CD25^+^FoxP3^+^ cells. C) Percentage of T-bet^+^CD4^+^ cells. D) Histogram of T-bet on CD4^+^ cells. Data ± SEM are representative of four independent experiments producing the same result profile. * P<0.05, **P<0.01, ***P<0.001.

### PD-L1 inhibits IFN-γ *in vivo* and regulates lesion size and parasite load

Next, we followed the *L. amazonensis* infection of C57BL/6 and PD-L1^−/−^ mice in terms of lesions size and parasite loads, and saw that PD-L1^−/−^ mice had greater lesion control and lower parasite burdens in the infected footpad after 9 weeks (Fig. 7A and B). We also investigated the production of IFN-γ and the transcription factor T-bet in CD4^+^ T cells from lymph nodes draining from the lesion. Infected PD-L1^−/−^ mice had higher frequencies of CD3^+^CD4^+^T-bet^+^IFN-γ^+^ T cells when compared to the infected C57BL/6 mice (Fig. 7C and D), but we did not see a difference in the Treg (CD3^+^CD4^+^CD25^+^FoxP3^+^) population *in vivo* (Fig. 7E and F). Corroborating our data, we observed an increase in IFN-γ production in the supernatant of the infected footpad macerate of PD-L1^−/−^ mice (Fig. 8A), as well as a reduction in IL-4 in this same group (Fig. 8B) when compared with infected C57BL/6 mice, demonstrating the favoring of the Th1 response. We did not observe any differences regarding IL-17 production in both analyzed groups (Fig. 8C).

**Fig 7.**
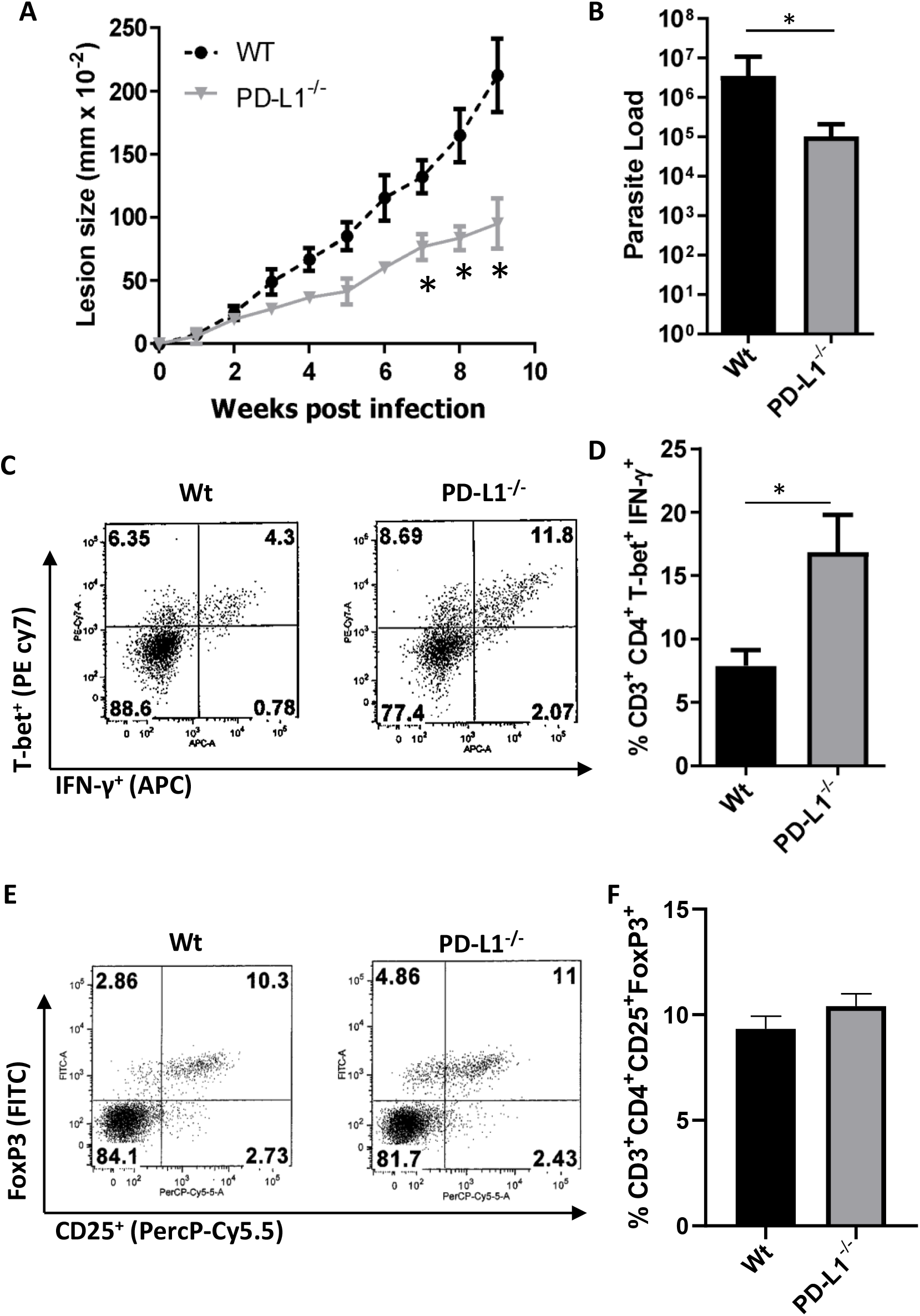
PD-L1 inhibits IFN-γ in vivo and regulates lesion size and parasite load. C57BL/6 (Wt) and PD-L1^−/−^ mice were infected in the hind footpad with L. amazonensis promastigotes (2×10^6^). After 9 weeks, the infected footpad and the draining lymph nodes were collected, macerated, and submitted to limiting dilution assay and flow cytometry, respectively. A) Lesion size as measured by digital caliper. B) Parasite loads from the infected footpads (5 mice/group). C) Dot plots of CD3^+^CD4^+^T-bet^+^IFN-γ^+^ cells (T-bet-PE-Cy7 and IFN-γ-APC). D) Percentage of CD3^+^CD4^+^T-bet^+^IFN-γ^+^ cells. E) Dot plots of CD3^+^CD4^+^CD25^+^ FoxP3^+^ cells (CD25-Percp-Cy5.5 and FoxP3-FITC). F) Percentage of CD3^+^CD4^+^CD25^+^FoxP3^+^ cells. Data ± SEM of individual mice (5 mice/group) are representative of four independent experiments producing the same result profile. * P<0.05.

**Fig. 8.**
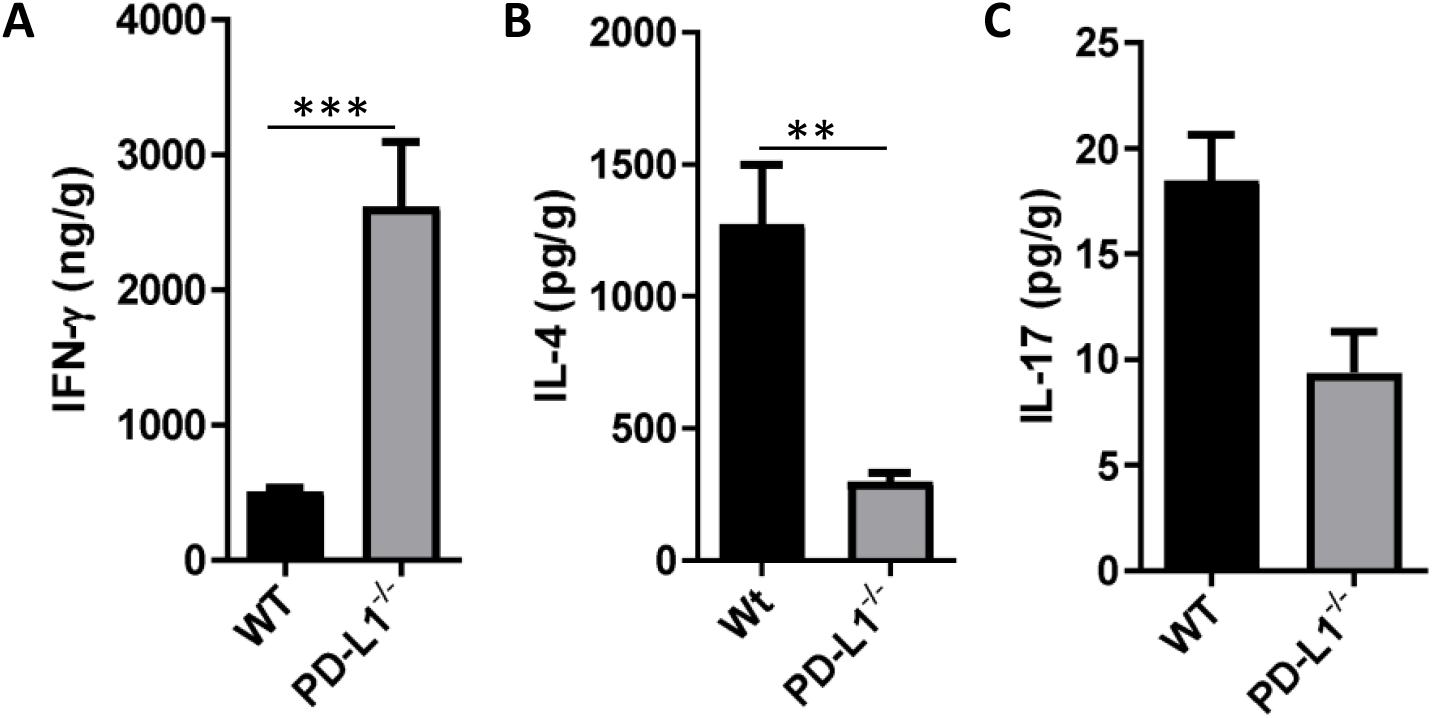
Increase of IFN-γ in the footpads of PD-L1^−/−^ mice infected by *L. amazonensis*. *L. amazonensis*-infected footpads were collected and macerated after approximately 9 weeks of infection and the supernatant was analyzed by ELISA. A) IFN-γ, B) IL-4, and C) IL-17. Data ± SEM of individual mice (5 mice/group) are representative of four independent experiments producing the same result profile. **P<0.01, ***P<0.001.

## Discussion

This study aimed to understand the induction of PD-L1 in the *in vitro* infection of DCs by *L. amazonensis*. In a previous work, we demonstrated that, when faced with *L. amazonensis* infection, neutrophils acquire a suppressor profile with increased PD-L1 expression in mouse ear lesions as well as in the draining lymph nodes. Furthermore, it was also observed that in co-culture assays of these PD-L1 expressing neutrophils with CD8 T cells there was a reduction in the production of IFN-γ, reinforcing the neutrophil suppressor role. Additionally, an increase in neutrophils expressing PD-L 1^+^ has been observed in lesions from patients infected with *L. braziliensis* (26).

DCs are the major regulators of the adaptive immune response as they educate T cells to assume either a T-cell activation phenotype or a T-cell tolerance phenotype (18). Tolerogenic APCs express PD-L1 on the surface, and *L. amazonensis* is able to induce the expression of PD-L1 on APCs.

There are different mechanisms to induce PD-L1 on DCs: secretion of IL-10 dependent on Toll-like receptors (TLRs), or the secretion of IL-10 and activation by JAK/STAT using PI3K-mTOR-STAT3 pathway or MAPK-ERK-STAT3 pathway. Cancer cells (29) and mesenchymal cells (27) produce IL-10 which activates STAT3, thereby promoting PD-L1 expression. *M. tuberculosis* (28) and HIV (29) have been reported to induce IL-10 production to promote PD-L1 expression. It is surprising that no increase in IL-10 production was detected from *L. amazonensis-*infected DCs (Suppl. Fig. 4B), but there was an increase when LPS was also incubated with the DCs (30). Furthermore, we did not observe any increase of PD-L1 expression on non-infected cells suggesting that there was no participation of cytokines in a paracrine manner, but we cannot exclude an autocrine mechanism. This observation indicates that *L. amazonensis* uses a different mechanism to induce PD-L1 expression.

*L. amazonensis* during the evolution process developed different pathways to drive the induction of tolerogenic APCs to suppress DC activation and prevent effector T cell differentiation. The PI3K-mTOR-STAT3 is known to enhance the viability and the survival of cancer cells (31). In leishmaniasis, since *L. mexicana* has been shown to inhibit apoptosis in DCs (32), species from the *L. mexicana* complex (including *L. mexicana* and *L. amazonensis*) could also use this pathway to induce PD-L1. The small participation of MyD88 in the induction of PD-L1 by *L. amazonensis* observed in the present study is very surprising. *M. tuberculsosis* (21), HIV (33, 34) and myeloma (27) are dependent on the TLR-MYD88 pathway for the induction of PD-L1 expression. We consider that the participation of MyD88 in our model may be in support of mTOR activation (35). Similar to cancer cells, *L. amazonensis* induces STAT3-PI3K-mTOR, however, the main mechanism is mTOR dependent.

Tolerogenic APCs are associated with the induction of Tregs *in vitro* and *in vivo* by the expression of PD-L1 (20). *M. tuberculosis* (21), *S. aureus* (22), and lung flora (23) promote Treg expansion via induction of PD-L1 on DCs. In a similar way, *L. amazonensis* induces Tregs *in vitro* (Fig. 6B).

We also observed an inhibition of the Th1 response in our model and, consequently, a reduction in the production of IFN-γ by T cells, corroborating previous work already mentioned (28). In another study, where we evaluated the treatment of leishmaniasis with anti-PD-1 and anti-PD-L1 monoclonal antibodies in a murine model of *L. amazonensis* infection, we observed an increase in the production of IFN-γ in CD4 and CD8 T cells, which consequently led to reduced parasite loads (11). The present study highlights the importance of PD-L1 since the observations of the previous work performed in the susceptible BALB/c mouse are reproduced in the partially resistant C57BL/6 mouse strain.

In summary, we could prove the participation of PD-L1 and PD-1 in inhibition of IFN-γ *in vitro* and *in vivo* by *L. amazonensis* infection and the ability of *L. amazonensis* to induce PD-L1 on DCs in the mechanism dependent of p-mTOR.

---

**Suppl. Fig 1: C57BL/6 mice infected with *L. amazonensis.*** C57BL/6 mice were infected in the footpad with *L. amazonensis* promastigotes (5×10^6^). Lesion size was measured weekly by digital caliper. After 3 months, footpads and draining lymph nodes were collected and macerated for the analysis of parasite load by limiting dilution assay and cellular composition by flow cytometry, respectively. Naïve = not infected. A) Lesion size. B) Parasite load of the infected footpad. C) Total number of total cells in the draining lymph nodes. D) Percentage of CD11c^+^ cells. E) Number of CD11c^+^ cells. F) Percentage of CD4^+^ T cells. G) Number of CD4^+^ T cells. Data ± SEM of individual mice (5 mice/group) are representative of three independent experiments producing the same result profile. * P<0.05, **P<0.01, ***P<0.001, ****P<0.001.

**Suppl. Fig 2: Induction of PD-L1 expression and inhibition of PD-L2 expression on infected dendritic cells.** BMDCs (5×10^5^) from BALB/c mice were infected for 18 h with amastigotes (Am; ratio 5:1) and promastigotes (Pro; ratio 10:1) of *L. amazonensis*. Medium = non-infected cells. Cells were analyzed by flow cytometry for CFSE^+^ (*L. amazonensis*), CD11c^+^ (PerCP-Cy5.5), PD-L1^+^ (APC), and PD-L2^+^ (PE). A) Percentage of CD11c^+^PD-L1^+^ cells. B) Percentage of CD11c^+^PD-L2^+^ cells. D) Percentage increase of PD-L1 expression. Data ± SEM are representative of three independent experiments producing the same result profile. *P<0.05, **P<0.01.

**Suppl. Fig 3: Western blotting analysis of p-mTOR and mTOR.** BMDC were infected or not with amastigotes (ratio 5:1) of *L. amazonensis,* overnight. p-mTOR and mTOR levels were analyzed by Western blotting. Glyceraldehyde 3-phosphate dehydrogenase (GAPDH) was used as an internal control.

**Suppl. Fig. 4: TGF-β and IL-10 levels in the supernatants of dendritic cell cultures.** BMDCs (5×10^5^) from BALB/c mice were infected for 18 h with *L. amazonensis* amastigotes (Am; ratio 1:5 or 1:20). LPS (100 ng/ml) was used as a positive control. Medium = non-infected cells. The supernatants were collected and analyzed by ELISA. A) TGF-β, B) IL-10. Data ± SEM are representative of three independent experiments producing the same result profile.

