## Supplementary figures and images for "*Leishmania amazonensis* infection induces PD-L1 expression on dendritic cells in an mTor-dependent manner and impairs Th1 responses *in vitro* and *in vivo*"

### Supplemnetary figures

**Supplemental Figure 1**

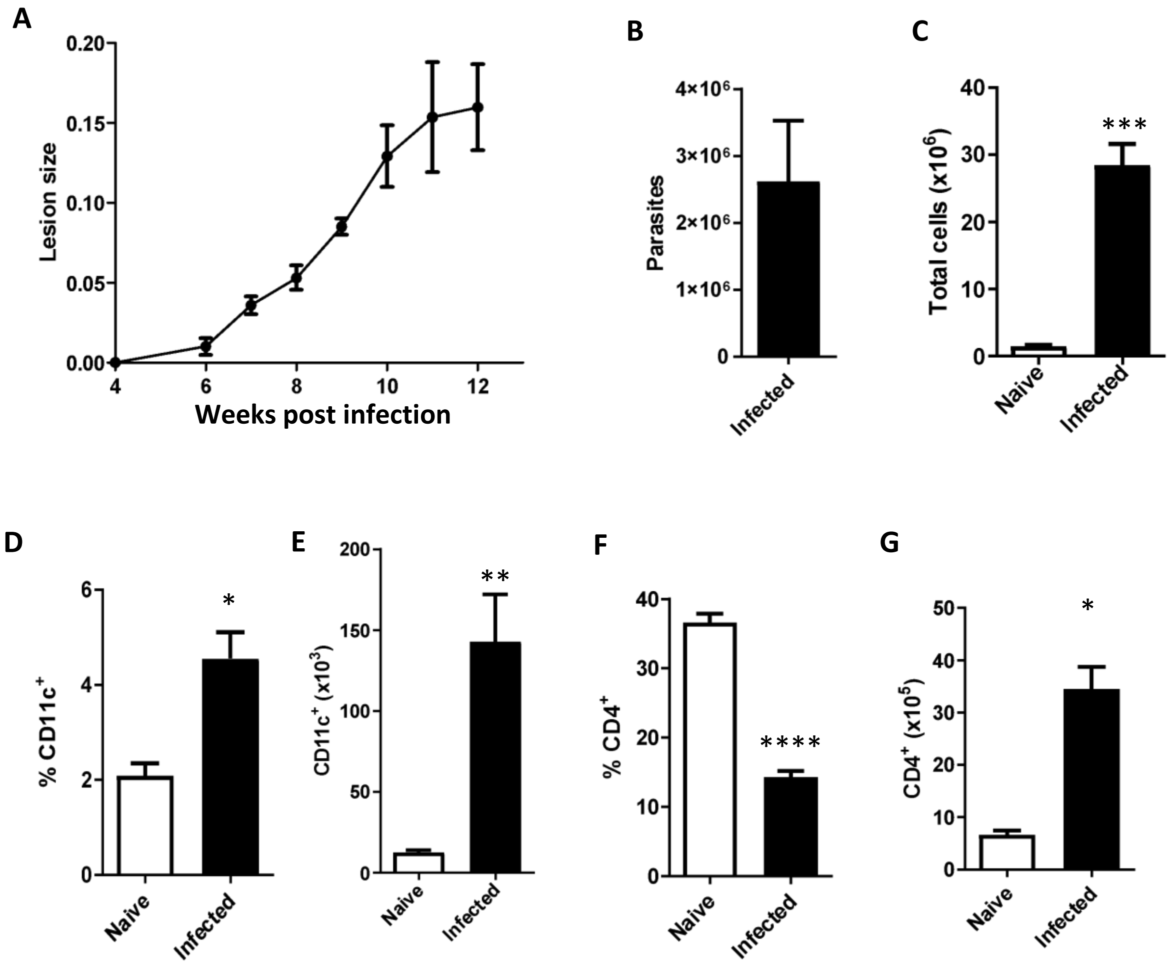

**Supplemental Figure 2**

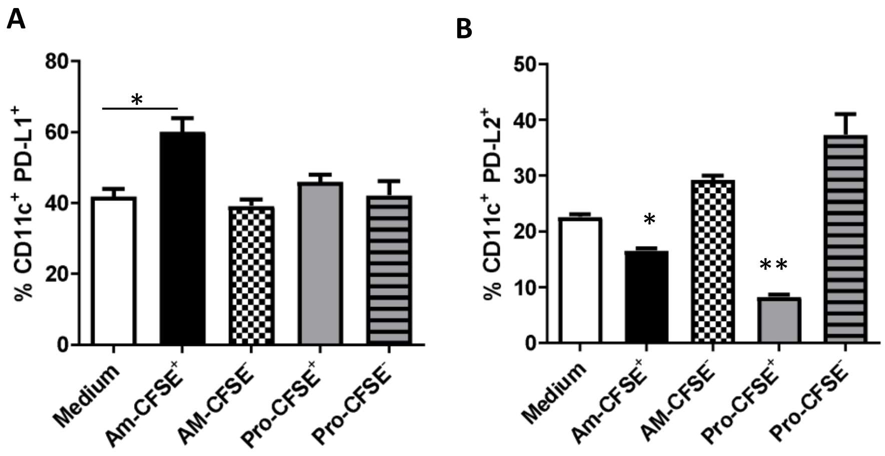

**Supplemental Figure 3**

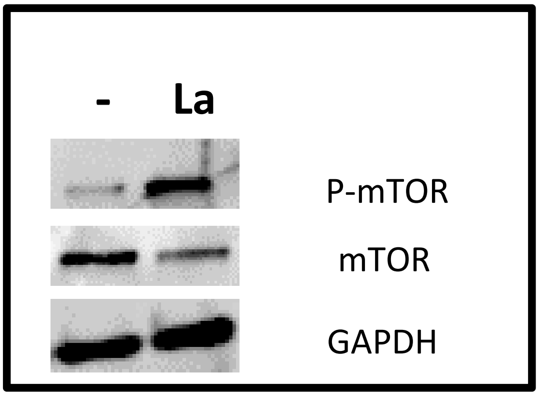

**Supplemental Figure 4**

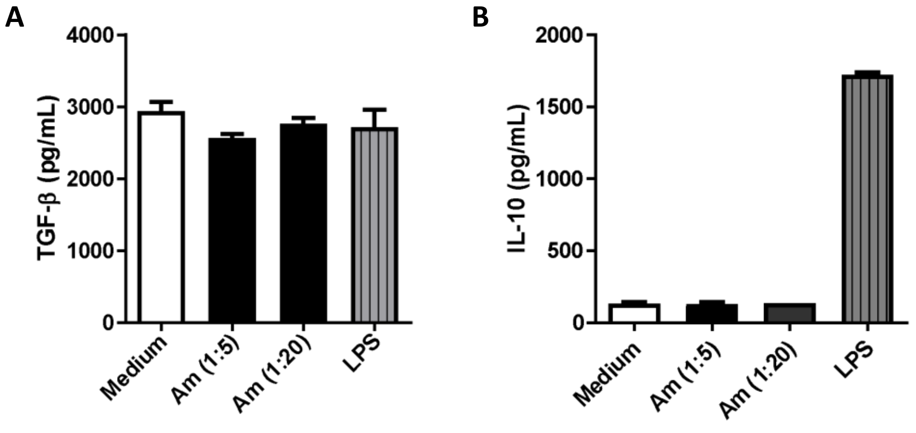
